# Eg5 Inhibitor SB743921 Causes p53-Dependent Cell Cycle Arrest, Senescence and Death in Tumor Cells

**DOI:** 10.1101/2025.01.23.634373

**Authors:** Iaroslav E. Abramenko, Alvina I. Khamidullina, Tamara A. Kiryukhina, Anna V. Tvorogova, Nataliya G. Pavlenko, Alexandra V. Bruter, Victor V. Tatarskiy

**Affiliations:** Laboratory of Molecular Oncobiology, Institute of Gene Biology, Russian Academy of Sciences, Moscow 119334, Russia; Center for Precision Genome Editing and Genetic Technologies for Biomedicine, Institute of Gene Biology, Russian Academy of Sciences, Moscow 119334, Russia

**Keywords:** SB743921, Eg5 inhibitor, Ixabepilone, mitotic inhibitors, p53, CKI, cell death

## Abstract

Mitotic inhibitors, such as Vinca alkaloids and taxanes, are one of the most effective chemotherapeutic agents used in the clinic. Despite their advantages, there are drawbacks to their use – primarily development of resistance and a high rate of side-effects, including damage to non-proliferating tissues. A range of new inhibitors targeting mitosis, whose activity does not depend on the binding to tubulin, are currently tested in clinical trials. Among such agents, inhibitors of Eg5 kinesin are highly promising due to their high activity and specificity. Here we show that compared to other drugs that target mitosis, an Eg5 inhibitor, SB743921, preferentially eliminates *TP53*-mutated cells and induces irreversible senescence, even after the drug washout, regardless of the p53 status. These effects are not defined by the immediate block of mitosis where SB743921 and a clinically used mitotic inhibitor Ixabepilone induce similar rates of mitotic arrest, apoptosis and induction of p53 and p21, but rather a long-term reaction, with absence of proteins required for replication, such as Cyclin A, E2F1, pRB. While after the washout Ixabepilone-treated cells can exit senescence and resume proliferation, cells treated with SB743921 did not exit the senescent state and did not resume proliferation as based on SA-β-galactosidase staining and EdU incorporation. The remaining senescent cells were effectively eliminated by Bcl2/Bcl-xL/Bcl-w inhibitor ABT-263, showing a potential of the combinational therapy with senolytic drugs. In total, we show the capacity of Eg5-targeting drugs for therapy of high-risk *TP53*-mutated tumors, which are potentially resistant to clinically approved mitotic inhibitors.

## Introduction

From 1961 when FDA approved vinblastine, mitotic microtubule inhibitors such as Vinca alkaloids (block tubulin polymerization) and taxanes (prevent depolymerization) became a backbone of chemotherapy, transforming oncology (1). They remain a mainstay of clinical care with new anti-mitotic drugs, such as epothilones being introduced into practice. However, there are many limitations of such treatments, mainly off-target toxicity and resistance through senescence, mitotic slippage and reversible mitotic arrest (2–4).

The status of the *TP53* gene is crucial for resistance to taxanes and Vinca alkaloids (5,6). The p53 transcriptional factor controls key pathways such as DNA damage response, cell cycle control and programmed cell death. More than 50% tumors contain mutations in *TP53* gene, which emphasizes a critical importance of the p53 protein in oncogenesis and tumor therapy (7,8). Overall, upregulated proliferation rate, metastasis and invasion activity, drug resistance and general aggressiveness are common traits of *TP53*-deficient tumor cells (9). These factors highlight the importance of searching for new effective agents against such cells.

Novel antimitotic drugs targeting Aurora kinases, Polo-like kinases, and mitotic kinesins are of great interest for cancer treatment (10). Inhibitors of mitotic kinesin motors are considered prospective due to their mechanism of action. Inhibition of KSP (kinesin spindle protein, Eg5) results in formation of monopolar mitotic spindles because of the inability of spindle microtubules with attached chromosomes to change their positions (11). To date, dozens of clinical trials with KSP inhibitors have been conducted. However, the majority of these drugs have not passed further than I/II phases (12). Their efficiency is limited by poor pharmacokinetics, insufficient information regarding mechanisms of action and tumor markers which will indicate best response, compared to other microtubule-binding agents. SB743921 was discovered alongside a number of Eg5 inhibitors, demonstrating high efficiency in preclinical models and indicating a favorable pharmacokinetic profile, with a half-live of 29 hours. SB743921 is one of the most selective Eg5 inhibitors with 40,000 fold selectivity against other kinesins (13). Recent works verified efficacy of SB743921 *in vivo* and revealed more detailed information about the cell death mechanisms after the drug exposure (14). Phase I trials also proved promising the potential of SB743921 but showed the necessity of proper dosage and delivery (15). Additionally, a recent publication of Li and colleagues exhibited a new target delivery method of SB743921 using antibody-drug conjugates (16).

In the present study we demonstrated the mechanism which determines an irreversibility of cell cycle arrest by SB743921 in tumor cell lines depending on the *TP53* status. Induction of the senescence is dependent on p53, and cells lacking p53 preferentially die by apoptosis. Furthermore, despite the similarities of primary molecular responses, long-term effects of Eg5 kinesin inhibitor and other antimicrotubule agents differ dramatically. SB743921 significantly lowers protein levels of E2F1, pRb, and Cyclin A leading to a prolonged arrest as well as to sensitivity to the senolytic ABT-263. Our work demonstrates the potential of new KSP inhibitors in selected tumors with mutated or deleted p53.

## Materials and Methods

### Cell culture

A549 (non-small cell lung cancer), HCT116 (colon cancer), MCF7 (breast cancer) cell lines were purchased from the American Type Culture Collection (ATCC) and cultured in DMEM (PanEco) supplemented with 10% fetal bovine serum (Biosera), 2 mM L-glutamine, 100 U/ml penicillin and 100 μg/ml streptomycin (PanEco) at 37°C, 5% CO_2_.

### Drugs

The following drugs were used: KSP (Eg5) inhibitor SB743921 (SB, Medchem Express), Ixabepilone (Ixa, Swords laboratories), Paclitaxel (Pacl, Fujian South Pharmaceutical), Vincristine (Vinc, Teva), Nocodazole (Noc, Calbiochem), MDM2 inhibitor Nutlin-3 (SelleckChem), Bcl-2/Bcl-xL/Bcl-w inhibitor ABT-263 (ApexBio) and Doxorubicin (SelleckChem).

### CRISPR/Cas9 mediated generation of TP53-/- cell lines

The plasmid based on the PX458 vector with gRNA sequence on the 2nd exon of *TP53* (CGACGCTAGGATCTGACTG) and the sequence encoding Cas9-GFP were used for transfection of the A549, HCT116 and MCF7 cells using GenJect 39 (Molecta). After overnight incubation with the vector, GFP-positive cells were sorted out using BD FACSAriaTM II system (BD Biosciences). Then cells were cultured for 3 days in fresh medium followed by the addition of Nutlin-3 (10 uM) for the following 3 days. These manipulations allowed the population to be enriched with cells carrying non-functional p53 and then subcloned to select for homozygous knockouts. Monoclonal colonies were examined by immunoblotting before and after p53 induction using 500 nM Doxorubicin for 4 hours (Supplementary Fig. S1).

### SRB assay

All assays were performed in 96-well plates (SPL Life Sciences). 1×10^3^ cells were added to each well, and on the next day mitotic inhibitors were added in the indicated concentrations for 7 days. Control cells were treated with equal concentrations of DMSO. After the incubation, media was removed, and samples in each well had been fixed with 10% TCA (trichloroacetic acid, NeoFroxx) for 10 min at +4°C. Wells were washed 3 times with dH2O, and then 0.4% SRB (Macklin) solution in 1% acetic acid was added for 10 min at room temperature. After that each well was washed with 1% acetic acid 5 times, and left overnight for full drying. Then SRB was dissolved in Tris-Base solution, pH 10, and had been incubated for 2 hours on orbital shaker OS-20 (BioSan). Optical density of samples was measured at 570 nm wavelength using CLARIOstar® Plus (BMG LABTECH).

### Colony formation assay

Cells were seeded onto 60 mm Petri dishes (SPL Life Sciences) at two different densities (5x10^2^/cm^2^ or 5x10^3^/cm^2^) and exposed to indicated compounds for 24 or 72 h (1d or 3d). Two densities were applied in order to compare the impact of cell density to the effectiveness of the drugs. After 24 or 72 h cells were washed twice in fresh medium for 15 min and further incubated in fresh media. After incubation cells were washed with PBS, fixed with methanol for 10 min at 4°C, and then stained with crystal violet solution (ChemCruz) for 10 min at room temperature, washed in water and dried overnight. Dishes were photographed using iBright FL1500 Imaging System (Invitrogen).

### Flow cytometry

Cells were plated at the density of 400x10^3^ cells in 6 cm dishes (19x10^3^/cm^2^) and treated with the indicated concentrations of drugs. After the completion of incubations, cells were lysed in a solution containing 50 μg/mL propidium iodide (Sigma-Aldrich), 100 μg/mL RNase A, 0.1% sodium citrate, and 0.3% NP-40 (VWR Life Science) and immediately analyzed on a Cytoflex Flow Cytometer 26 (Beckman Coulter). At least 10,000 fluorescent ‘events’ were collected per each sample. For data processing, the CytExpert Software (Beckman Coulter) and GraphPad Prism 9 software were used.

### Reverse transcription and qPCR

Cells were plated at the density of 400x10^3^ cells in 6 cm dishes (19x10^3^/cm^2^). Total RNA was isolated using the RNeasy Mini Kit (QIAGEN). RNA concentrations were measured on the spectrophotometer Implen Nanophotometer P300 (Implen). Reverse transcription was performed applying MMLV reverse transcriptase, mQ water, DDT, dNTP mix, OligodT primer (Evrogen), 5х Reaction buffer (Thermo Scientific) and isolated RNA. cDNA samples were synthesized using the aforementioned mix and were additionally dissolved in 80 μL mQ water to achieve 5-fold dilution. For qPCR in real time we used 5x qPCRmix-HS, EvaGreen 20x fluorescent dye, mQ water and primers. All reagents were purchased from Evrogen. qPCR were performed on LightCycler 96 (Roche) using the following mode: Denaturation: 95°С, 60 sec; 39 cycles of amplification: 95°С 20 sec, 62°С 19 sec, 72°С 15 sec; Termination and melting: 95°С 10 sec, 65°С 60 sec, 97°С 1 sec.

The *RPLP0* transcript was used for signal normalization. Each experiment contained negative controls with RNA and premixes without cDNA. Primer sequences are presented in Supplementary Table S1.

### Immunoblotting

Cells were lysed in 100-200 μL solution containing 50 mM Tris-HCl pH 8.0, 150 mМ NaCl, 0,1% sodium dodecyl sulfate, 1% NP-40, 2 mМ phenylmethylsulfonyl fluoride (VWR Life Science) and protease inhibitor cocktail, (Sigma-Aldrich) for 30 min on ice. Total protein concentration was measured by the Bradford method. Proteins were resolved by electrophoresis in 10-12% polyacrylamide gels (30-50 μg protein per lane) and transferred on 0.2 µm nitrocellulose membranes (BioRad). Non-specific protein-protein interactions were blocked with 5% skimmed milk for 30 min at room temperature. Membranes were incubated with primary antibodies (Supplementary Table S2) overnight at 4°C and washed three times in TBS-Tween-20 (1:1000). Secondary antibodies conjugated with horseradish peroxidase were diluted in 5% skimmed milk in TBS and incubated for 1 h at room temperature. Then membranes were washed three times in Tween-20-TBS (1:1000). Bands were visualized using Clarity Western ECL Substrate (Bio-Rad) on a iBright FL1500 Imaging System.

### EdU staining and confocal microscopy

On the indicated day after a drug removal (7d), 5-ethynyl-2′-deoxyuridine (EdU, 10 µM) was added for 2 h at 37°C. Cells were fixed with 4% paraformaldehyde (Penta Chemicals) solution in PBS for 15 min, gently washed with PBS and permeabilized in 0.2% solution of Triton X-100 in PBS for 30 min and washed with PBS. Freshly prepared mixture of 2 mM Cu(II)-BTTAA, 5 µM AF488 azide and 10 mМ ascorbic acid (Lumiprobe) in 100 mМ PBS in 50% DMSO was added for 30 min. After that, cells were washed in PBS, stained with DAPI (Thermo Fisher, 5 µg/µL in PBS) for 10 min and mounted in Mowiol 4-88 (Sigma-Aldrich). Detection of fluorescence was performed on a Leica TCS SP2 confocal microscope (Leica Microsystems). Analysis of images was performed using ImageJ software.

### Senescence-associated β-galactosidase detection

Cell samples after the indicated treatment were fixed in a solution containing 2% paraformaldehyde and 0.2% glutaraldehyde in PBS for 5 min at room temperature. Then cells were washed with PBS and exposed to the staining solution containing 1 mМ MgCl_2_, 3.3 мМ K_3_Fe(CN)_6_, 3.3, mМ K_4_Fe(CN)_6_, 0.02% NP-40 and 0.04% Х-Gal (Helicon) pH 6.0 for 16 h at 37°С. Images were captured on a Nikon Eclipse TI microscope (Nikon).

It is important to note that as negative controls we used untreated cells and cells treated with non-toxic concentration of ABT-263. For these controls cells were seeded at lower density and incubated for a shorter period (4 days) to avoid overgrowth.

### Statistics

Comparisons were performed using a one-way ANOVA test with a multiple comparisons Dunnett’s test or a two-way ANOVA test with a multiple comparisons Sidak’s test (GraphPad Prism 9.0). Data are presented as mean ± standard deviation of at least 2 independent biological replicates. The P value < 0.05 (*) was taken as evidence of statistical significance.

## Results

### 1) SB743921 is more potent than conventional mitotic inhibitors and selectively eliminates A549 *TP53* knockout cells

We treated MCF7, A549 and HCT116 cell lines with the microtubule-stabilizing agents Ixabepilone (Ixa) and Paclitaxel (Pacl), microtubule-destabilizing compounds Nocodazole (Noc) and Vincristine (Vinc) or KSP inhibitor SB743921 (SB) for 7 days followed by SRB tests to compare the cytotoxic potencies of different mitotic inhibitors (Fig. 1A, Supplementary Figure S2).

**Figure 1.**
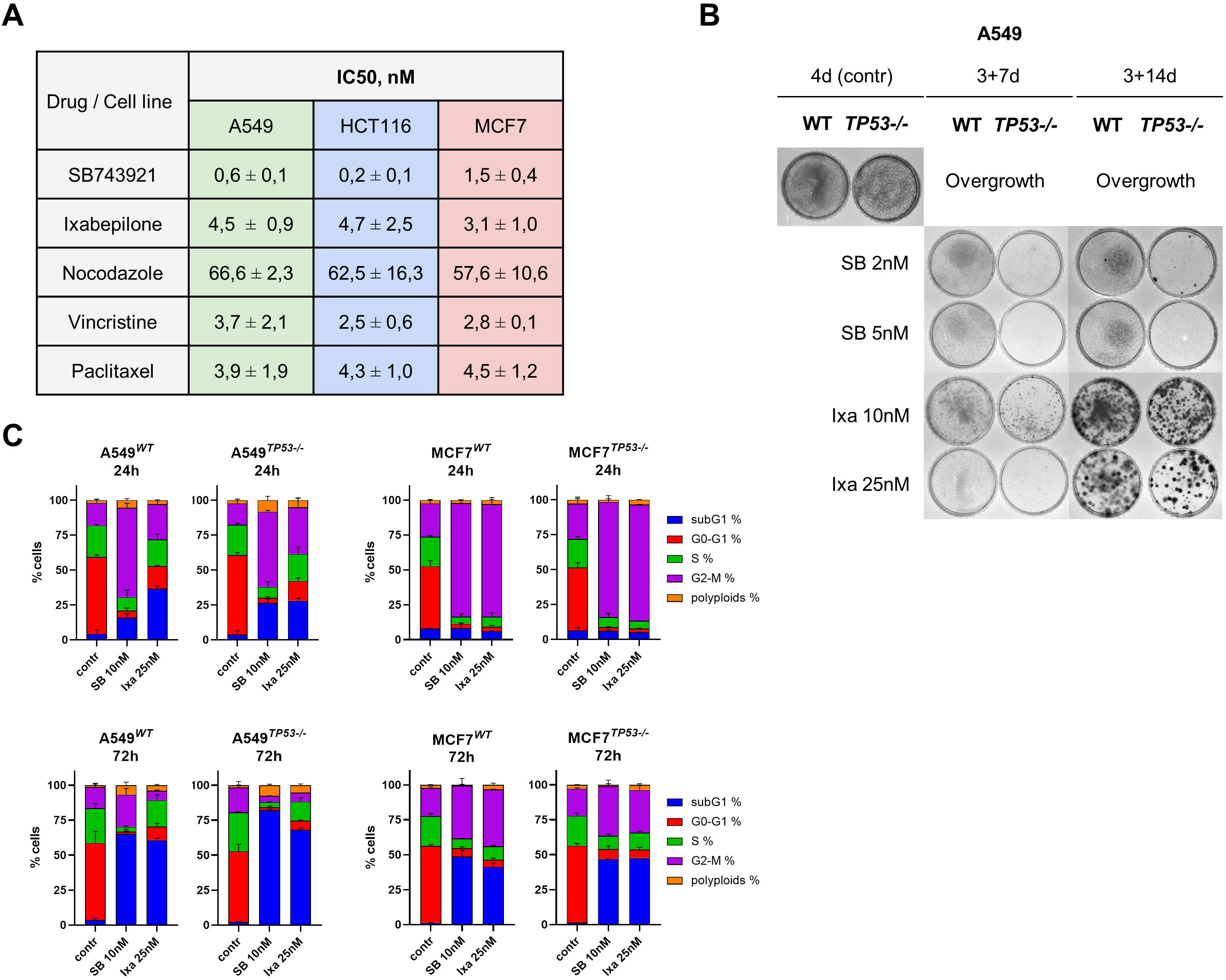
Cytotoxicity of SB743921 against mitotic inhibitors in cells with the wild type (WT) and knocked out p53 (*TP53-/-*). **A**, IC50 values (mean ± SD, n=3) of A549, MCF7 and HCT116 exposed to various mitotic inhibitors for 7 days were obtained based on the SRB assay. **B**, Proliferative and colony formation ability of A549*^WT^* and A549*^TP53-/-^* after 7 d or 14 d from the removal of SB743921 (SB) or Ixabepilone (Ixa). Cells were exposed to drugs for 72 h and then incubated in fresh media the following 7 d or 14 d. **C,** Cell cycle distribution of A549*^WT^* and A549*^TP53-/-^* (*left panels*) and MCF7*^WT^* and MCF7*^TP53-/-^* (*right panels*) treated with SB (10 nM) or Ixa (25 nM) for 24 h (*upper panels*) and 72 h (*bottom panels*).

As shown in Fig. 1A and S2, Noc was the least effective mitotic inhibitor; Ixa, Pacl and Vinc demonstrated intermediate values, while SB was the most potent. Efficacy of other microtubule poisons did not depend on the cell line, while SB was more effective against HCT116 (IC_50_ **≈** 0.2 nM) and less effective against MCF7 (IC_50_ **≈** 1.5 nM). In all tested cell lines, SB performed as the most effective mitotic inhibitor. To observe long-term effects of tested drugs on cell survival, we studied colony formation by A549, MCF7 and HCT116 cells after treatment with SB or Ixa (Fig. 1B and Supplementary Fig. S3). Ixa caused relatively long-lasting arrest in all tested cell lines; however, there were colonies that resumed growth after 2 weeks (Fig. 1B and Supplementary Fig. S3). In contrast to this, SB-treated A549 cells did not resume growth even after 14 days after the drug washout (Fig. 1B). Surprisingly, the A549*^TP53-/-^* subline was even more affected, compared to p53 proficient cells, with almost no cells surviving. Therefore, SB can be effective for elimination of p53-deficient cells.

Cell density can affect the response to cytotoxic drugs, because quiescent cells can escape from the mitotic block and secrete pro-survival factors in conditioned media. We performed colony formation assay with a sparse cell density to examine the duration and reversibility of different mitotic inhibitors to test the impact of mentioned factors (Supplementary Fig. S4). This experiment clearly demonstrated that Ixa, Pacl and Vinc have durable but reversible effects, while SB caused an extremely prolonged antiproliferative effect even by 2 weeks after the drug removal. In SB-treated samples, very few colonies were noticed by day 21 post washing of a low drug concentration (2 nM), indicating a remarkable duration of SB antiproliferative effect. Thus, SB performed as the most prolonged mitotic inhibitor indicating that this compound has a distinct mechanism of action.

To examine the mechanism of prolonged SB’s anti-proliferative effects and increased cytotoxicity in p53-negative cells, we decided to compare it with one of the clinically used tubulin-binding agents – Ixa. First we compared cell cycle distribution for both drugs in equitoxic concentrations. Both SB and Ixa, as expected, caused G2/M arrest in MCF7 cells after 24 h incubation (Fig. 1C, *upper right*). However, the proportion of A549 cells arrested in G2/M was markedly bigger if cells were incubated with SB than with Ixa. Ixa did not cause a significant G2/M block after 24 h but instead increased the subG1 fraction of cells with fragmented DNA (Fig. 1С, *upper left*). Additionally, at this time point, we did not observe a difference between cells with wild type *TP53* and *TP53-/-*. After 72 h incubation, MCF7 cells appeared to be less responsive to SB and Ixa than A549 based on the subG1 fraction (Fig. 1С, *bottom panels*). Interestingly, there was no significant difference in cell cycle distribution in response to these compounds, nor was there any difference between MCF7*^WT^*and MCF7*^TP53-/-^* (Fig. 1С, *right panels*). In addition, the A549^TP53-/-^ subline was more sensitive to SB, as indicated by the subG1 population, compared to A549^WT^ cells although the difference was insignificant (P value = 0.075). This effect was further substantiated by the SRB test with A549*^WT^* and A549*^TP53-/-^*, which proved higher sensitivity of A549*^TP53-/-^* cells to SB (Supplementary Fig. S4).

### 2) SB and Ixa induce similar short-term responses of cell cycle arrest and induction of apoptosis but different long-term effects

To investigate the difference in response to SB or Ixa that lead to the striking differences in the longevity of their effects (Fig. 1B), we examined the expression of proteins that regulate cell cycle transition and cell death of MCF7 and A549 cell lines after 24 h (Fig. 2A). Both mitotic inhibitors caused an increase of p53 and p53-dependent cyclin-dependent kinase inhibitor (CKI) p21. Meanwhile, the p53-independent CKI, p18, was not changed in response to SB or Ixa. Nevertheless, *TP53-/-* cells were also arrested in G2/M, demonstrating that p53 is not required for induction of mitotic arrest. Additionally, we observed an increased cleavage of poly (ADP-ribose) polymerase (PARP1), a hallmark of apoptotic cell death, in response to SB and Ixa. Together, this data shows similarity between relatively short-term responses to SB and Ixa in A549*^WT^* and A549*^TP53-/-^* cells, suggesting that the prolonged anti-proliferative effect of SB does not depend on the short-term response to the drug.

**Figure 2.**
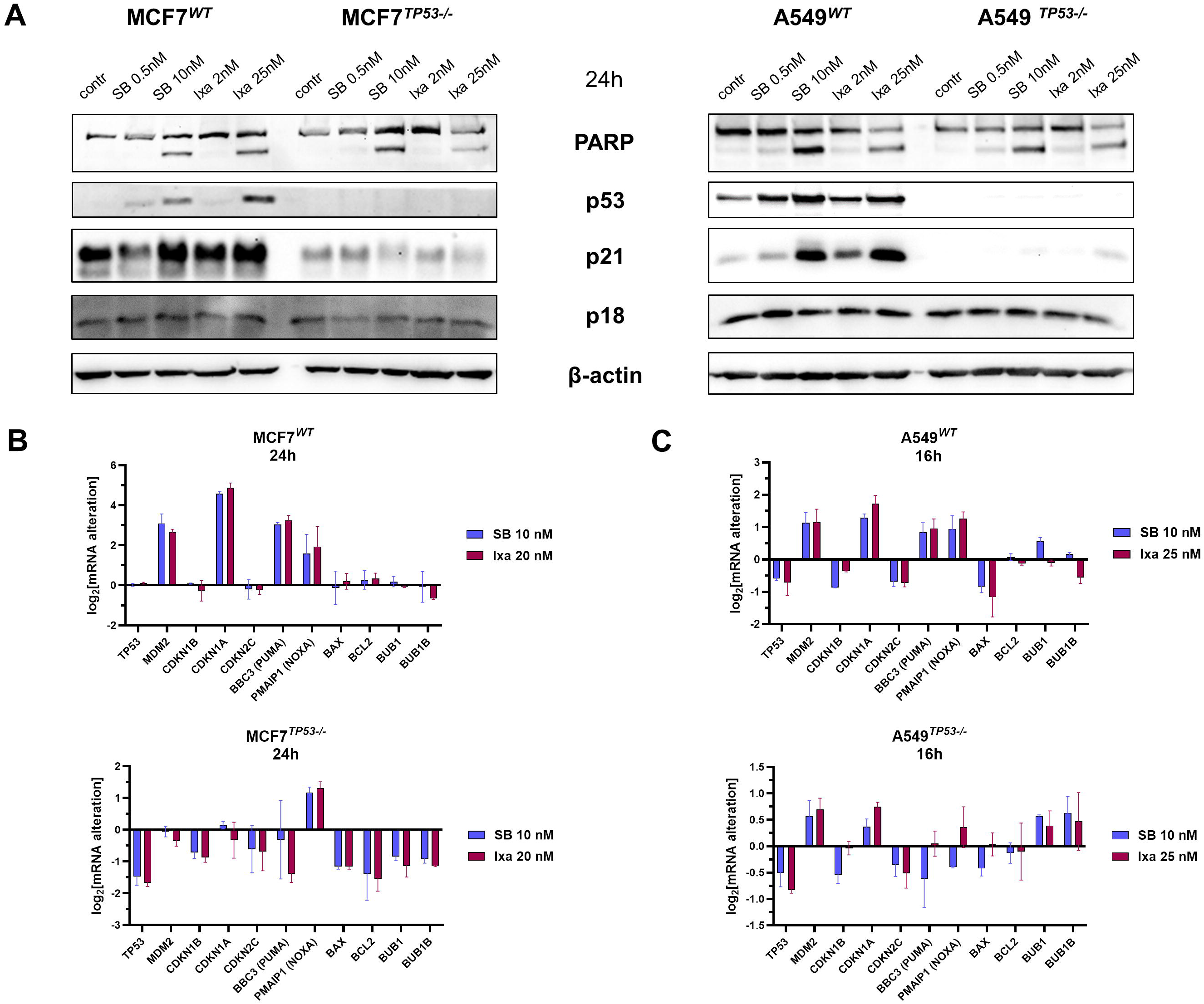
p53-mediated cell cycle inhibitory and proapoptotic pathways are activated in response to the exposure of SB743921 (SB) or Ixabepilone (Ixa). **A,** Immunoblotting analysis of MCF7 and A549 treated with SB or Ixa in different concentrations (SB 0.5 nM and 10 nM, Ixa 2 nM and 25 nM) for 24 h. **B, C**, qPCR analysis of MCF7 and A549 respectively treated with SB (10 nM for MCF7, 3 nM for A549) or Ixa (25 nM) for 16 h (A549) or 24 h (MCF7).

Gene expression analysis using qPCR confirmed the induction of the p53-dependent *CDKN1A* gene encoding CKI p21 and the absence of activation of the *CDKN2C* and *CDKN1B* genes that encode p53-independent CKIs p18 and p27 (Fig. 2B and C). The transcripts of p53-dependent pro-apoptotic PUMA and NOXA (*BBC3* and *PMAIP*) were also similarly upregulated by SB or Ixa. The *BCL2L2* gene, which encodes the p53-independent antiapoptotic protein Bcl-w, and the p53-dependent proapoptotic *BAX* were not changed considerably in cells with the wild type *TP53*. These results are in agreement with the necessity of p53 for induction of PUMA and NOXA, but not BAX (8). Expression of the *MDM2* gene, a negative p53 regulator, increased in response to SB and Ixa. In general, a noticeable difference between *WT* and *TP53-/-* cells consisted of the induction of p53- dependent cell cycle inhibition (p21) and proapoptotic pathways (PUMA, NOXA) in cells with the wild type *TP53* that was not observed in *TP53-/-.* Overall, quantitative alterations in protein and mRNA levels were distinct in A549 and MCF7, however, a general pattern of changes was similar for both cell lines, considering their diverse origin. The most remarkable difference was observed only between *TP53WT* and *TP53-/-* cell cultures.

Investigations of protein and mRNA levels after 72 h were complicated by low numbers of surviving cells, so further experiments were performed with drug washout and cell cultivation in fresh medium. This allowed us to analyze long-term effects of SB or Ixa in surviving cells.

### 3) Prolonged action of SB may be explained by irreversible cell cycle arrest and vast cell death, especially in A549*^TP53-/-^* cells

As there was no apparent difference in mechanisms of cell death and cell cycle arrest (Fig. 2), which could explain the difference in long-term efficiency of SB and other mitotic inhibitors in colony assays (Fig. 1B, Supplementary Fig. S4), we examined delayed effects of SB and Ixa. Flow cytometry analysis after 3 days and 6 days after the drugs removal (Fig. 3A and B) showed that after treatment with Ixa for 24 h followed by washing off the drug, cells re-entered the cycle and resumed proliferation 6 days post treatment (Fig. 3B). The most noticeable effect was a higher ratio of cells in the S phase compared to untreated control cells (Supplementary Fig. S6).

**Figure 3.**
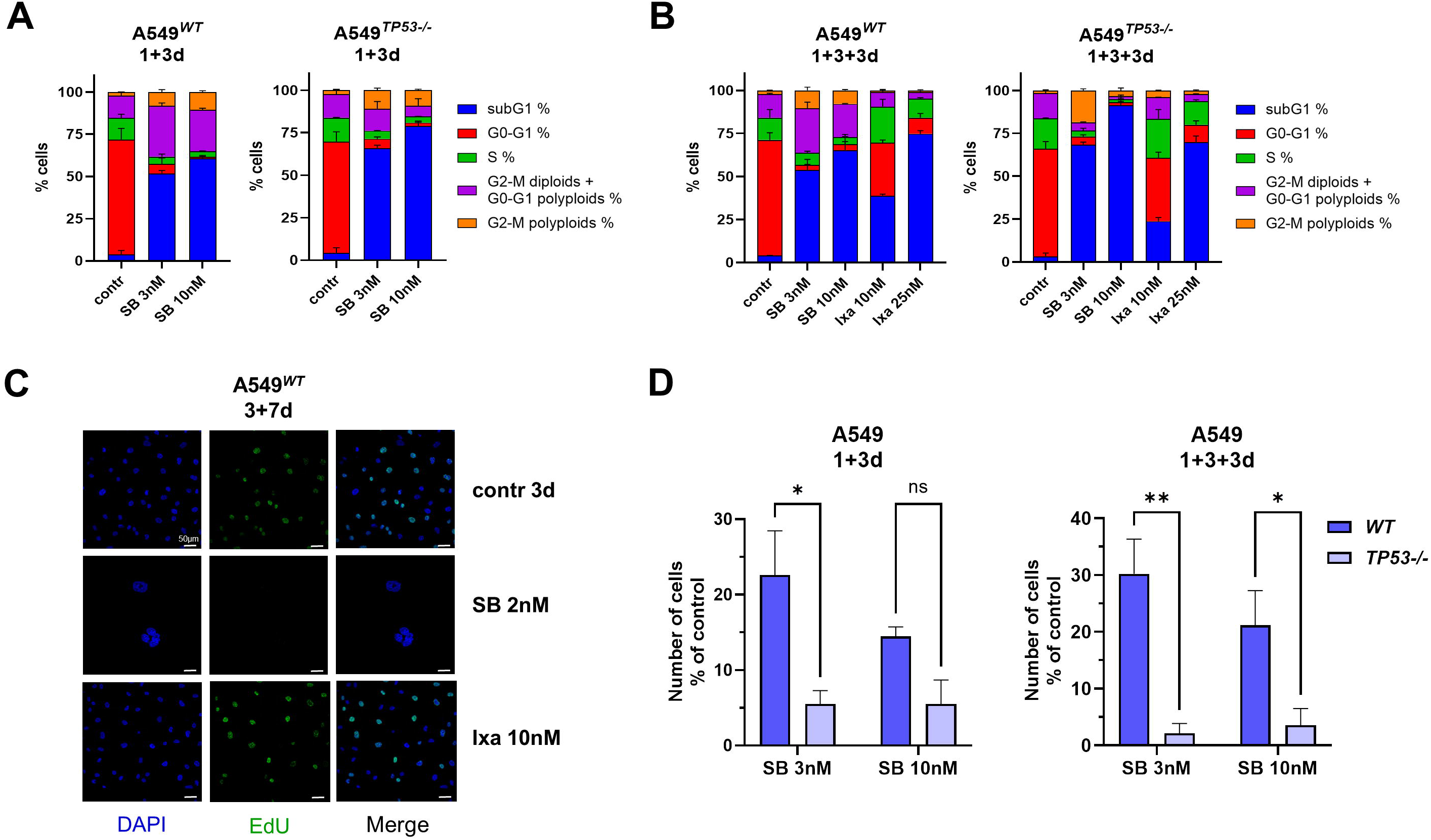
SB743921 induces irreversible proliferative arrest and predominant death of A549*^TP53-/-^*cells. **A**, **B**, Cell cycle distribution after 6 days after removal of SB743921 (SB) or Ixabepilone (Ixa). Cells were exposed to the drugs for 24 h, washed and incubated for 3 or 6 d in fresh media. **C**, Confocal microscopy images of EdU-treated samples after a week since the removal of SB or Ixa. Cells were exposed to drugs for 72 h before the washout. **D**, Single cell concentrations in flow cytometry analysis as a percentage of the untreated control samples. *, P < 0.05, **, P < 0.01; ns – not significant.

Contrary to the effect of Ixa, A549 treated with SB demonstrated a significant dependence on the p53 status. SB treatment in A549*^WT^*cells led to the accumulation of 4n cells (Fig. 3A and B). As was previously shown, one of the major mechanisms which leads to the survival after mitotic arrest is mitotic slippage with further polyploidization (3). Therefore, we hypothesized that A549*^WT^* were arrested in G2/M diploid and G1 polyploid fraction after the release from mitotic arrest. Based on EdU incorporation in surviving cells (Fig. 3C, Supplementary Fig. S7), A549*^WT^* did not undergo the subsequent replication (Supplementary Fig. S7 and S8). As a result, the majority of surviving cells were arrested in the post-slippage 4n G1/S. On the contrary, A549*^TP53-/-^* tended to resume replication after polyploidization. These cells, however, were not able to enter mitosis and continue their division that is evidenced by an accumulation of 8n tetraploid cells in the G2/M phase (Supplementary Fig. S8 and S9). In contrast to SB, we observed DNA synthesis in Ixa-treated A549*^WT^* which indicates a progression of replication (Fig. 3C, Supplementary Fig. S7). Additionally, SB-treated cells contained giant nuclei that were not found in Ixa-treated samples.

Overall, the survival rate of A549*^TP53-/-^*, compared to *WT*, was significantly lower after 6 days in drug-free medium (Fig. 3D). It can be associated with A549*^TP53-/-^*tendency to enter replication and therefore die. Thus, after G2/M arrest and subsequent polyploidization, A549 cells with functional p53 are arrested on the boundary of G1/S, whereas p53-null cells either die or undergo polyploidization and become arrested in 4n G2/M after the completion of replication. The sequence of events depending on the *TP53* status is presented in Supplementary Fig. S9.

### 4) Replicative signaling is abrogated in SB-treated cells

To reveal why SB demonstrated a sustainable inhibitory effect on the cell cycle, we investigated the levels of CKIs p18, p21, p27 after drug washout and long-term incubation (Fig. 4A). In A549*^WT^* cells, SB elevated p53 and p21 levels, while decreased p18. Similar effects were caused by Ixa, except for p18 which was less attenuated. Levels of p27 were unaltered by SB or Ixa. Other CKI proteins are not present in A549 according to the analysis of the DepMap database (https://depmap.org/portal/) and existing literature that shows that A549 cell line carries deletions of *CDKN2A* and *CDKN2B* and decreased mRNA synthesis of *CDKN1C* gene, which encode CKIs p14, p16, p15 and p57, respectively (17–19). Hence, cell cycle control 6 days after the SB or Ixa removal does not depend on the majority of CKIs – p14, p15 p16, p18 and p27. Thus, in the A549 line p21 can be considered a key regulator of cell cycle inhibition after the treatment with SB and Ixa. It is also important to note that p21 is the only CKI which is directly regulated in a p53-dependent manner (20,21).

**Figure 4.**
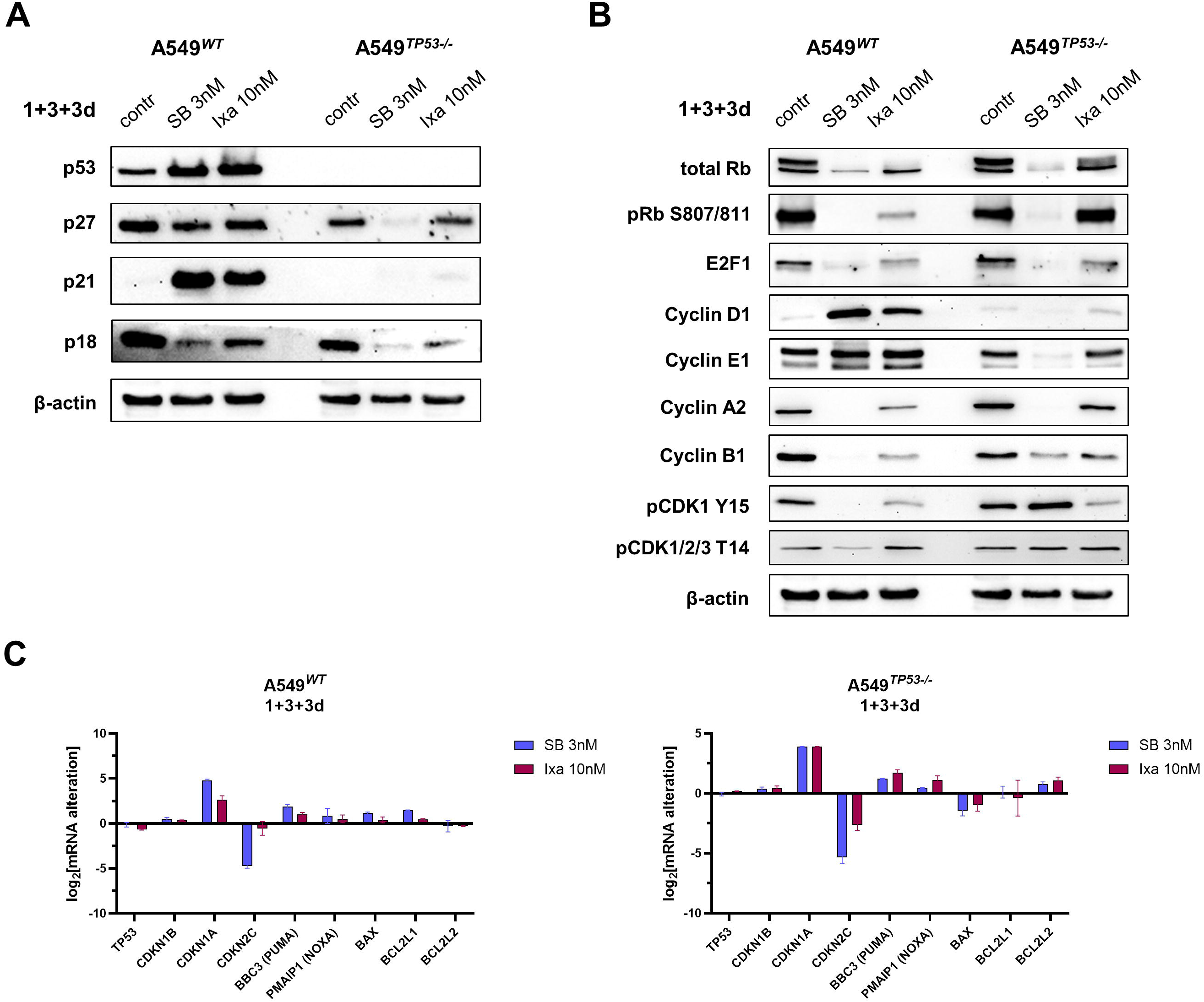
SB743921 depletes pro-replicative signaling causing irreversible arrest, while p53 is crucial for G1/S arrest in *WT* cells. **A**, Immunoblotting analysis in A549*^WT^* and A549*^TP53-/-^* cells 6 days after removal of SB743921 (SB) or Ixabepilone (Ixa). Cells were exposed to drugs for 24 h, then washed and incubated for 6 d in fresh media. **C**, qPCR analysis of A549*^WT^* and A549*^TP53-/-^* expression after 6 days since the removal of SB or Ixa. Cells were exposed to drugs for 24 h before the washout.

In the A549*^TP53-/-^* subline, in which p53 and p21 were not detectable (Fig. 4A) the level of p18 also decreased after the treatment with SB or Ixa. Interestingly, the p27 content substantially decreased in response to SB, but not to Ixa. No CKIs upregulation in *TP53-/-* cells evidenced that CKI do not prevent these cells from entering and undergoing replication, but nevertheless there is a mechanism that prevents further cell divisions.

Next, we evaluated mRNA levels changes after the drugs washout (Fig. 4C). Similarly to the changes on the protein level, in *WT* cells there was a considerable increase of the *CDKN1A* mRNA in SB-treated cells and a lesser, but still noticeable, elevation in response to Ixa. This effect, as well as a significant decline of *CDKN2C* in SB-treated cells, correlates with protein level of the respective mRNAs. However, *CDKN2C* mRNA in Ixa- treated *WT* cells and *CDKN1B* mRNA in SB-treated *TP53-/-* cells were unchanged, whereas at the protein level we observed a decline of these gene products, which indicates the existence of an additional regulation. It can also be confirmed by the *CDKN1A* upregulated expression in response to SB and Ixa in A549*^TP53-/-^*, as we observed no p21 accumulation. Alterations in mRNA levels of the apoptosis-related proteins was insignificant, with the exception for more than threefold elevation of p53-dependent *BBC3* (PUMA) in A549*^WT^*.

As demonstrated above, immediate cell cycle arrest by SB and Ixa was not significantly distinct (Fig. 4A and C), but we observed significant differences in the longevity of antiproliferative effects (Fig. 3A, D, and F). As there were no significant differences in cell cycle inhibitory proteins that could explain the difference between SB and Ixa, we next examined cell cycle promoting factors. Protein analysis showed that in A549*^WT^*, in 6 days after Ixa washout there was no considerable effect on pro-replicative factors, indicating a partial return to proliferation. In striking contrast, treatment of A549*^WT^* with SB led to a near total decline of major replication proteins pRb, E2F1, and Cyclins A2 and B1 (Fig. 4B). At the same time, the level of Cyclin E1 was unchanged, whereas Cyclin D1 dramatically increased compared to untreated cells. Together with the ploidy data (Supplementary Fig. S8), this indicated that SB-treated cells were arrested at the G1/S boundary.

In A549*^TP53-/-^*cells, Ixa also caused little effect on the replication markers E2F1, Cyclin A2, and pRb after washout (Fig. 4B). Effects of SB on pRB, E2F1 and Cyclin A2 levels in A549*^TP53-/^*were similar to A549*^WT^* cells, with complete depletion of these proteins. On the other hand, in A549*^TP53-/-^*the levels of Cyclins D1 and E1 decreased in SB-treated samples indicating the absence of G1/S-arrested cells. Instead, SB-treated A549*^TP53-/-^*cells expressed Cyclin B1, a marker of G2/M block, which was previously confirmed by the cell cycle analysis (Fig. 3B, Supplementary Fig. S9). At the same time, upregulation of the CDK1 phosphorylation in response to SB explains G2/M arrest after relatively successful completion of replication. This inhibitory phosphorylation of CDK1 by Wee1 may be a major mechanism explaining the inability of A549*^TP53-/-^*cells to undergo the next round of division after SB-induced mitotic arrest even after 6 days in drug-free medium (Supplementary Fig. S9).

### 5) SB induces irreversible senescence in A549 cells: sensitivity to senolytics

As SB-treated A549*^WT^* cells did not undergo replication and expressed replication proteins, we hypothesized that these cells were irreversibly arrested and underwent senescence. To confirm this we stained cells for senescence-associated β-galactosidase (SA- β-Gal, Fig. 5A). Up to 93% of A549*^WT^* cells were senescent after 6 days in the drug-free medium whereas treatment with Ixa did not cause vast senescence in the surviving population (Fig. 5A and B). A549*^TP53-/-^* cells were less prone to undergo senescence (Fig. 5B): only ∼43% cells were senescent after treatment with SB. Nevertheless, despite the fact that A549*^TP53-/-^* lack the markers of senescence such as p53, p21 and p16, these cells were also stained positively. We hypothesized that this effect is mediated through prolonged arrest of A549*^TP53-/-^* in G2/M in response to SB, for which p53, p21 and p16 are not mandatory.

**Figure 5.**
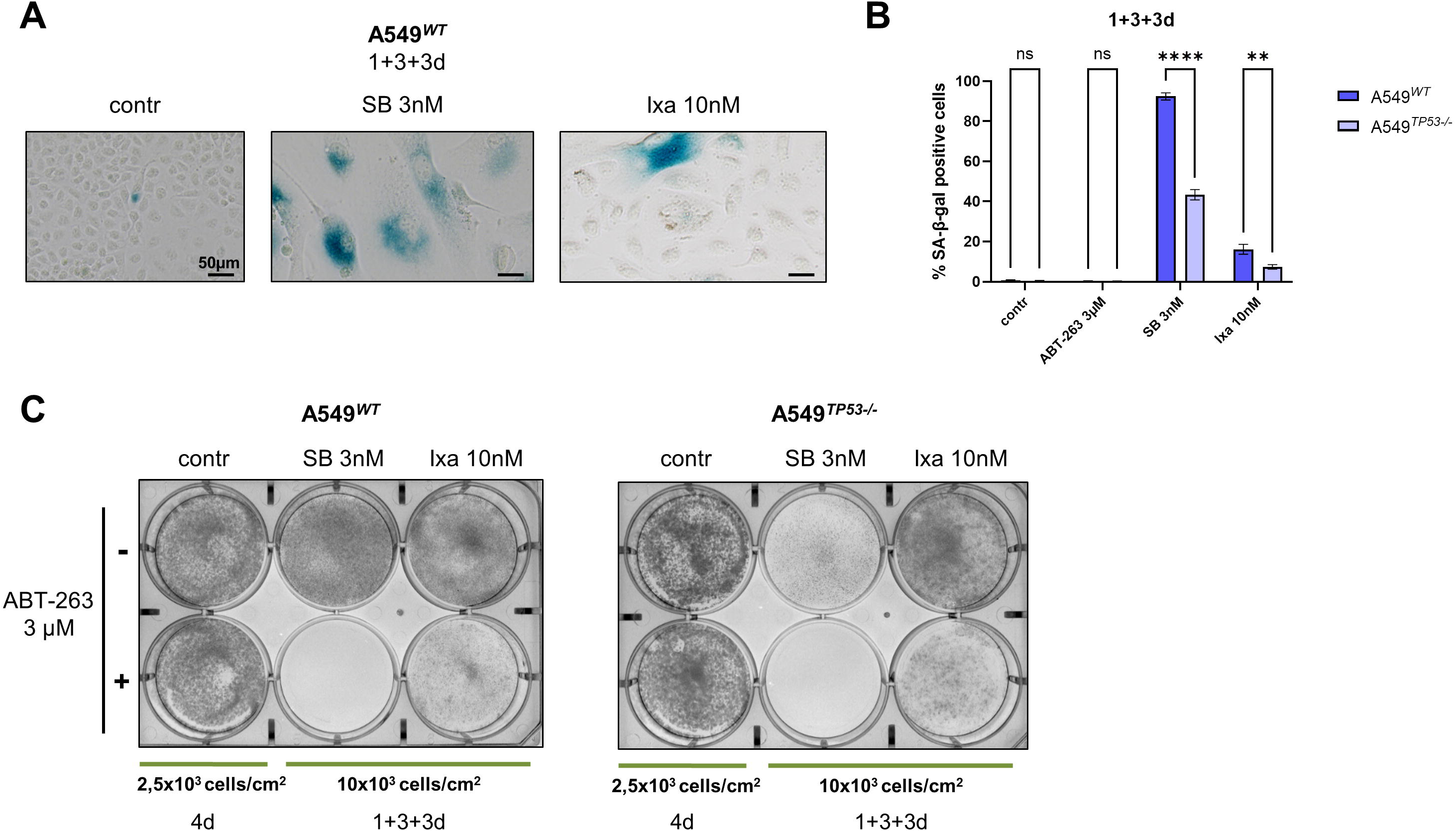
SB743921 induces irreversible senescence in A549*^WT^* and SB743921-treated senescent cells can be targeted by senolytics, such as ABT-263. **A**, Light microscopy images of A549*^WT^* 6 days after the removal of SB743921 (SB) or Ixabepilone (Ixa). Cells were fixed and then stained with solution for the SA-β-Gal detection. **B,** Quantitative analysis of stained A549*^WT^* and A549*^TP53-/-^* cells as demonstrated in **A**. Data were compared using the ANOVA test; **, P < 0.01, ****, P < 0.0001; ns – not significant. **C,** Crystal violet staining of A549*^WT^* and A549*^TP53-/-^* treated with SB or Ixa alone or together with 3 uM ABT-263.

As a result, the majority of A549*^WT^* remained in senescent condition after the SB treatment. Hence, these cells were not eliminated as effectively as A549*^TP53-/-^*, but at the same time they were unable to proliferate. However, dormant cells may be potentially dangerous due to the acquisition of senescence-associated secretory phenotype (22,23).

Therefore, we tested combinations with the senolytic compound ABT-263 (24) added after the removal of SB or Ixa (Fig. 5C). We used a non-toxic concentration (3 µM) which was sufficient for elimination of senescent cells formed upon the treatment with SB. The combination of ABT-263 and Ixa was less efficient because of a significantly lower ratio of senescent cells in Ixa-treated samples (Fig. 5B). Thus, we demonstrated the potential of combining the Eg5 inhibitor SB and the senolytic ABT-263 for elimination of p53-deficient cells and survived senescent *WT* cells.

## Discussion

Mitotic inhibitors, like taxanes and Vinca alkaloids are a mainstay of cancer therapy. Notwithstanding the high efficiency of tubulin-targeting drugs there are major drawbacks for their use. Mainly, these dose-limiting toxicities, including peripheral neuropathy and neutropenia (25), acquired resistance. One of major factors of resistance is deletion or mutation of *TP53*, a major tumor suppressor, which can lead to mitotic slippage (2) or escape from therapy-induced senescence (26,27).

But, despite our increasing understanding of regulation and mechanics of mitosis, new modalities of targeting mitotic cells, such as Aurora A/B inhibitors, PLK inhibitors and others, remain in the early stages of clinical trials. Eg5 inhibitors were considered as a prospective class of new-antimitotic drugs in the 2010s. Early work has indicated that they can overcome resistance to taxanes (28). Despite their high *in vitro* and *in vivo* activity, the clinical data for most of them proved insufficient for approval (29). One major drawback of the first generation of Eg5 inhibitors was their poor pharmacokinetics that led to insufficient half-lives for optimal dosing. New developments in the field with improved PK/PD properties, such as filanesib (30,31) and antibody-drug conjugates ((32), NCT06034275) maintain a probability for them entering clinical practice. An important part for adoption of targeted drugs is selection of markers which can predict the highest response rate. In this work we examined prospective indications where Eg5 inhibitors can prove to be more efficient than clinical tubulin-binding drugs in p53 mutated tumors. Recent data demonstrated that inhibitors of another mitotic kinesin, KIF18A, are more active in *TP53* mutated tumors (33).

First, we confirmed high efficiency of the Eg5 inhibitor SB743921 in three commonly used cell lines – HCT116, MCF7 and A549 (Fig. 1A, Supplementary Fig. S2). Surprisingly, in long term colony formation assays SB prevented appearance of colonies after the washout, especially in isogenic *TP53* knockout cells (Fig 1B). This difference was not present for tubulin-binding agents (Fig 1B, Supplementary Fig. S3), despite a very similar initial cell cycle arrest (Fig 1C). It is necessary to emphasize that the efficacy of utilized mitotic inhibitors greatly depends on the selected cell line. Unexpectedly, Ixa was more active against MCF*^TP53-/-^*, compared to MCF7*^WT^* (Supplementary Fig. S3), but this difference was leveled out after 2 weeks since the drug washout. In HCT116, Ixa caused a week-long arrest of proliferation as well, but after 14 days we observed many growing colonies (Supplementary Fig. S3). SB also demonstrated long-lasting antiproliferative activity against HCT116, and similarly to A549, HCT116*^TP53-/-^* were more sensitive to SB, compared to HCT116*^WT^*. Hence, we observed that sensitivity of A549 and HCT116 to SB relied on the p53 status, while MCF7 did not experience this effect. It can be explained by the difference in mechanisms by which these cell lines escape mitotic arrest. It is known that MCF7 cells contain micronuclei, and formation of this type of chromosomal instability commonly does not lead to the p53 activation (34,35), whereas A549 and HCT116 undergo micronucleation. It is also worth noting that the level of p53 did not differ substantially between A549, HCT116 and MCF7 (Supplementary Fig. S1), as also evidenced by approximately equal *TP53* expression levels (Supplementary Fig. S10, data is taken from the DepMap database).

Next, we examined the effects of the drugs on the cell cycle and the immediate response in expression of CKI proteins and apoptosis markers. We chose to focus on a comparison between SB with Ixa, a semi-synthetic analog of epothilone B (36), as one of the most recently approved tubulin-binding drugs. The immediate response to both drugs was very similar with G2/M arrest, subsequent apoptotic cell death (Fig 1C), similar rates of induction of p53, and cleavage of PARP1 (Fig. 2A). Notably in A549 Ixa induced more cell death than in MCF7, and a number of cells were able to proceed through G1 and S phases (Fig. 1C). Similarly the transcriptional response to these drugs was strikingly analogous (Fig. 2B and C). Nevertheless, after the drug washout Ixa-treated cells were able to resume proliferation with 78-87% of surviving cells moving through G1 and S phases after 6 days of media replacement (Fig. 3B) and a considerable proportion of cells incorporating EdU (30.7%) (Fig. 3C, Supplementary Fig. S7). On the contrary, SB-treated cells were arrested in G2/M and polyploid phases of cell cycle and were not able to resume replication, as evidenced by no EdU incorporation (Fig. 3C, Supplementary Fig. S7).

There are a number of mechanisms that prevent cells from resuming proliferation even after the washout of inhibitors. The activity of CDK-cyclin complexes can be blocked by the interaction with CKIs or by inhibitory phosphorylation by Wee1-family kinases (37).

We first examined the levels of CKI proteins that present in A549 (Fig. 4A). P21 was increased in both SB and Ixa-treated A549 cells 6 days after the washout, while p18 was downregulated by both SB and Ixa. In A549*^TP53-/^*^-^ cells, p21 was not induced on the protein level and p18 was even more downregulated. It is important to note the existence of a previously described p53-independent pathway of p21 expression (38), as we similarly observed upregulation of p21 mRNA levels, but not on the protein level (Fig. 4C). As other CKI are not present in A549 cells, and the levels of p21 and p53 were very similar after exposure to both SB and Ixa, we presumed that the resumption of proliferation is mediated by other mechanisms. Indeed, in SB-treated cells E2F1 and Cyclin A, which are required for replication were absent in both *TP53WT* and *TP53-/-* cells. Interestingly, Wee1-dependent phosphorylation of CDK1 was present and increased in *TP53-/-* cells, but only an extremely low level of CDK1 phosphorylation was present in *TP53WT* cells (Fig. 4B). We propose that the G1/S block in *TP53WT* cells is mediated by p21 and absence of E2F1, while in *TP53-/-* cells, where p21 is absent, the arrest at the G2/M boundary is maintained by Wee1 activity. Indeed, it is well established that Wee1 inhibition promotes premature entry into mitosis, leading to increased cell death (39). This could be considered as a further strategy for treatment of Eg5i-treated cells, together with senolytics.

We hypothesized that the complete absence of proteins which are presented in replication and required for its initiation is a marker of senescent phenotype in SB-treated cells. To validate the presence of this phenotype we stained cells for a common marker of senescence – SA-β-Gal. We observed that even 6 days after the drug washout SB-treated A549*^TP53WT^*cells were positive for this marker, while Ixa-treated cells mostly exited from senescence (Fig. 5A and B). In A549*^TP53-/-^* cells, fewer cells were stained with SA-β-Gal which corresponds with the known role of p53 in a maintenance of senescence (26,27). However, it should be mentioned that the SA-β-Gal detection assay cannot be accurate and specific to the necessary extent. Senescence poses various risks due to the acquisition of senescence-associated secretory phenotype, which may lead to a development of a drug resistance and occurrence of side effects (23). Additionally, there is existing evidence about so-called phenomena “neosis”, which means that some senescent cells can escape their dormant state and start to proliferate again after a long period because of the potential generation of small mononuclear cells (27,40). Thus, senescent cells can be dangerous in terms of tumor recurrence as well. A number of drugs, called senolytics, were shown to eliminate senescent cells *in vitro* and *in vivo* (*41*). Senescent cells express high levels of anti-apoptotic Bcl-family proteins, and Bcl-family inhibitors were shown to work as senolytics. We demonstrated that addition of Bcl2/Bcl-xL/Bcl-w inhibitor – ABT-263 – completely eliminates remaining SB-treated cells, while having no effect on control cells, and little effect on Ixa-treated cells (Fig. 5C). Such a strategy is considered for enhancing activity of mitotic inhibitors (10) and our data demonstrates that Eg5 inhibitors are a very promising partner for such combinations.

Overall, our data suggests that Eg5-targeting drugs can be used for tumors, where other mitotic inhibitors are not likely to work, especially in p53-mutated cancers. Eg5- targeted drugs are also a promising pairing for use with apoptosis-inducing and senolytics drugs such as Bcl-inhibitors. Future work should be done to validate markers for tumors, where Eg5 inhibitors could show the most benefit.

## Authors’ Contribution

**I. Abramenko:** Conceptualization, data curation, formal analysis, investigation, methodology, visualization, writing – original draft. **A. Khamidullina:** Data curation, formal analysis, investigation, visualization, writing – review and editing. **T. Kiryukhina:** Methodology. **A. Tvorogova:** Methodology. **N. Pavlenko:** Methodology. **A. Bruter:** Methodology. **V. Tatarskiy:** Conceptualization, resources, data curation, formal analysis, supervision, funding acquisition, validation, investigation, project administration, writing – original draft, review and editing.

## Acknowledgements

The authors are grateful to A.A. Shtil for reviewing the original draft and for valuable contribution to the project, I.A. Vorobjev and G.S. Kopeina for discussions, and D.M. Potashnikova for assistance in cell sorting.

## List of Abbreviations

CDK: cyclin-dependent kinase
CKI(s): cyclin-dependent kinase inhibitor(s)
EdU: 5-ethynyl-2′-deoxyuridine
Ixa: Ixabepilone
KSP: kinesin spindle protein
Noc: Nocodazole
Pacl: Paclitaxel
PARP1: poly (ADP-ribose) polymerase 1
SA-β-Gal: senescence-associated β-galactosidase
SB: SB743921
Vinc: Vincristine

