## Supplementary Fig. S for "Eg5 Inhibitor SB743921 Causes p53-Dependent Cell Cycle Arrest, Senescence and Death in Tumor Cells"

### Supplementary Materials
(10 figures)

#### Eg5 Inhibitor SB743921 Causes p53-Dependent Cell Cycle Arrest, Senescence and Death in Tumor CellsAuthors

Iaroslav E. Abramenko, Alvina I. Khamidullina[1,2], Tamara A. Kiryukhina[1], Anna V. Tvorogova[2], Nataliya G. Pavlenko[1,2], Alexandra V. Bruter[2], Victor V. Tatarskiy[1,2]

**Affiliations**

[1]Laboratory of Molecular Oncobiology, Institute of Gene Biology, Russian Academy of Sciences, Moscow 119334, Russia

[2]Center for Precision Genome Editing and Genetic Technologies for Biomedicine, Institute of Gene Biology, Russian Academy of Sciences, Moscow 119334, Russia

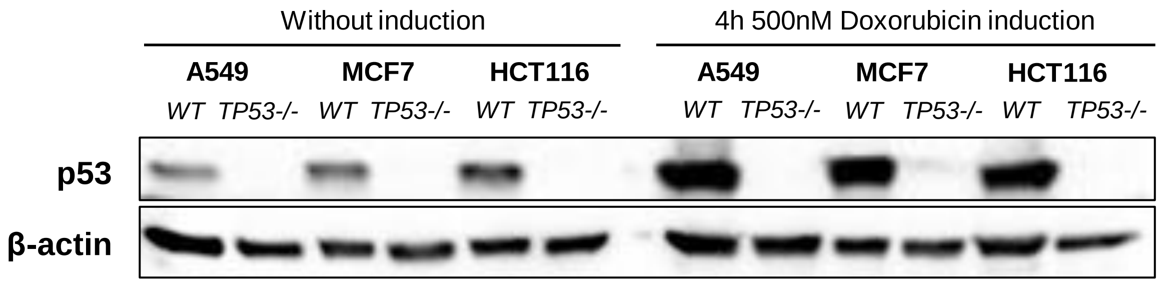


**Supplementary Figure S1.** Test of TP53 knockout in A549, MCF7, and HCT116 cell lines. Immunoblotting before and after p53 induction using 500 nM Doxorubicin for 4 hours.

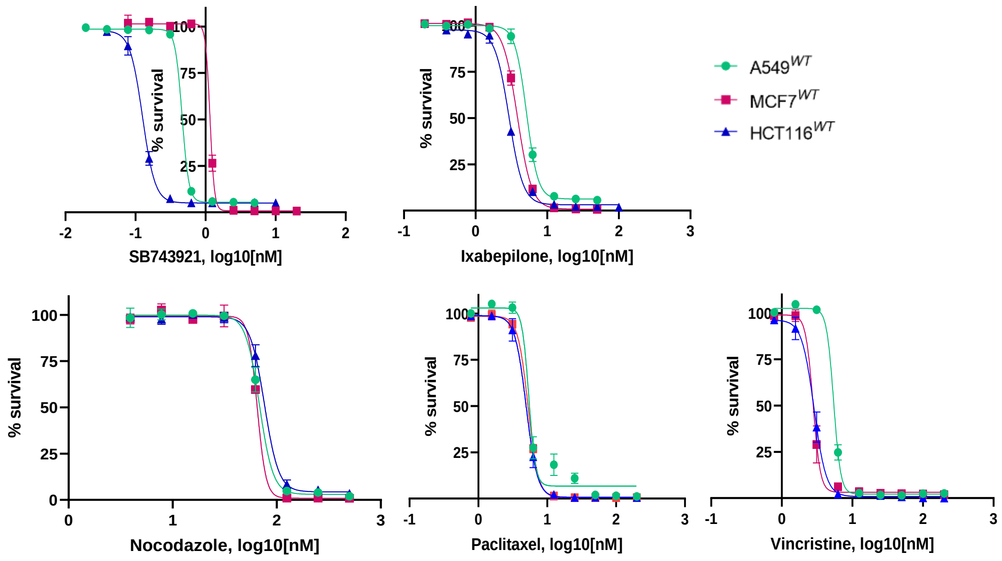


**Supplementary Figure S2.** Survival curves of A549, MCF7 and HCT116 exposed to various mitotic inhibitors for 7 days analyzed by SRB assay.

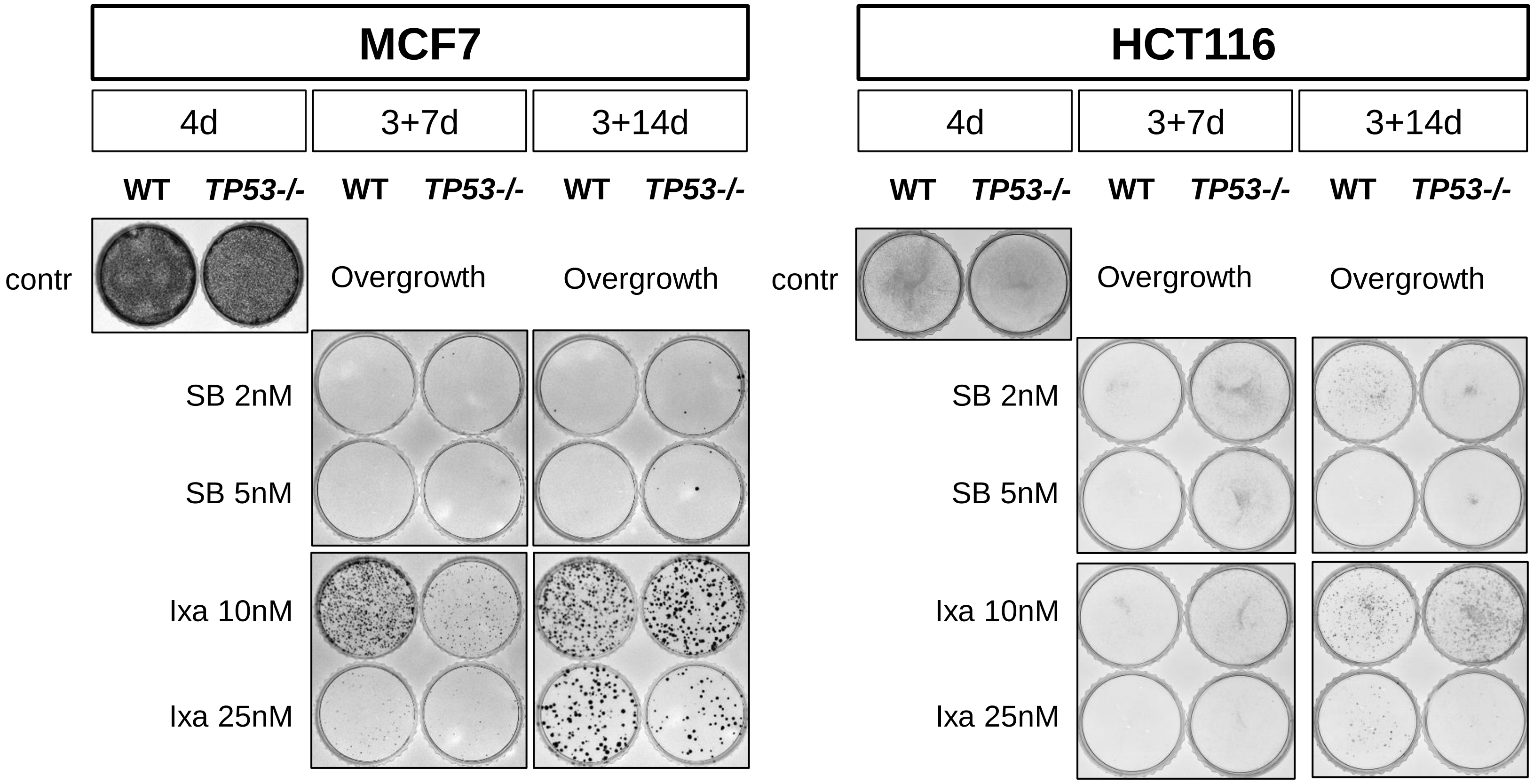


**Supplementary Figure S3.** Proliferative and colony formation ability of $MCF7^{WT}$, $MCF7^{TP53-/-}$ (*left panels*) and $HCT116^{WT}$, $HCT116^{TP53-/-}$ (*right panels*) after a week and two weeks since the removal of SB743921 (SB) or Ixabepilone (Ixa). Cells were exposed to the drugs for 72 hours before the washout.

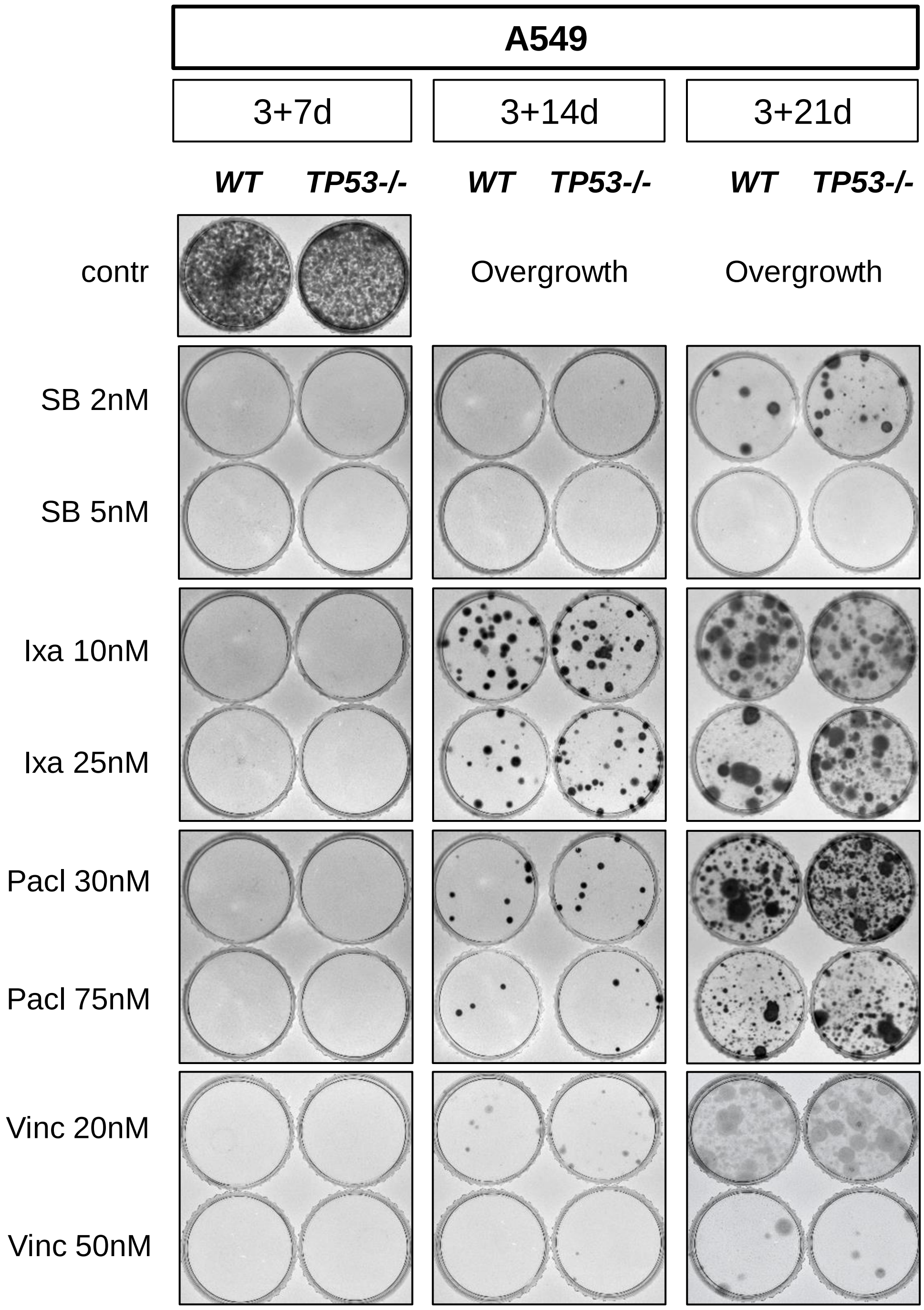


**Supplementary Figure S4.** Proliferative and colony formation ability of A549WT and A549TP53-/- after a week, two and three weeks since the removal of SB743921 (SB), Ixabepilone (Ixa), Paclitaxel (Pacl), and Vincristine (Vinc). Cells (in low density 5x102 cells/cm2) were exposed to the drugs for 72 hours before the washout.

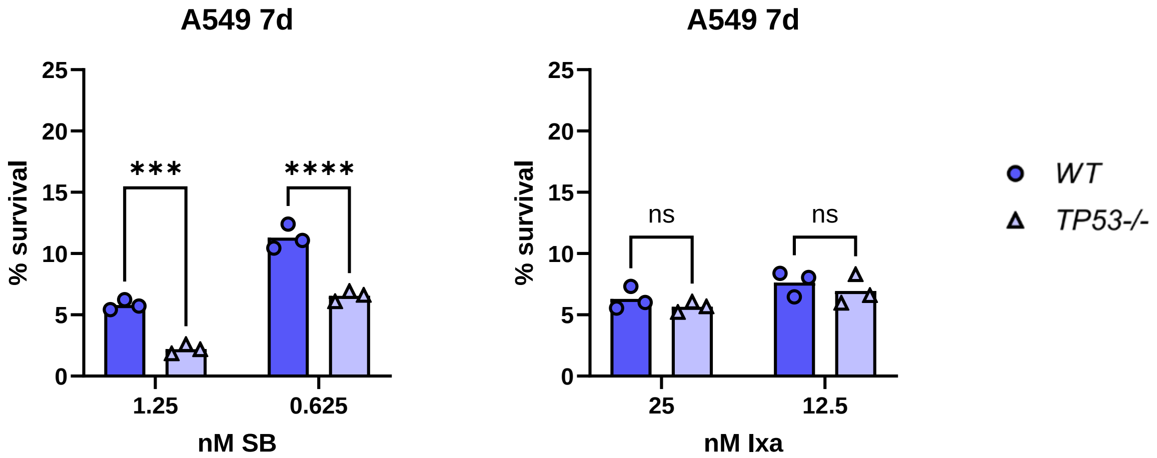


**Supplementary Figure S5.** $A549^{TP53-/-}$ was more sensitive to SB743921(SB)-induced cell death. SRB test with $A549^{WT}$ and $A549^{TP53-/-}$ after 7 days of 1.25nM and 0.625nM SB, 25nM and 12.5nM Ixabepilone (Ixa). Data are represented as the mean (n = 3). Data were compared using the ANOVA test; ***, $P < 0.001$ ; ****, $P < 0.0001$; ns – not significant

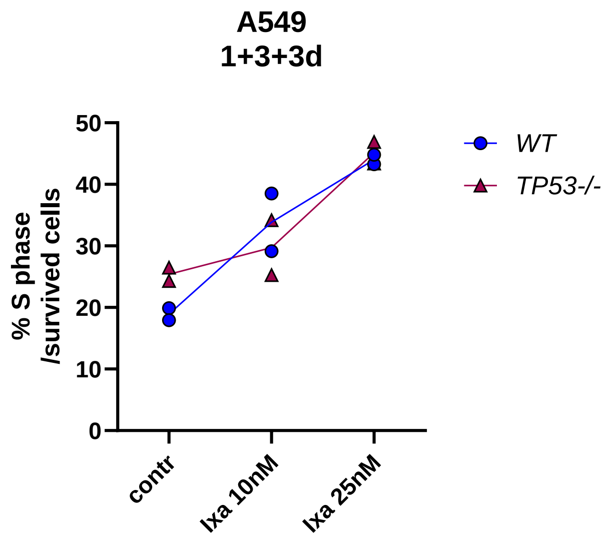


**Supplementary Figure S6.** Ratio of cells in S phase to all survived cells. A549$^{WT}$ and A549$^{TP53-/-}$ cells treated with 10 nM SB743921 (SB) or 25nM Ixabepilone (Ixa) for 1 day and subsequent 2 washouts every 3 days. Quantification based on flow cytometry data.

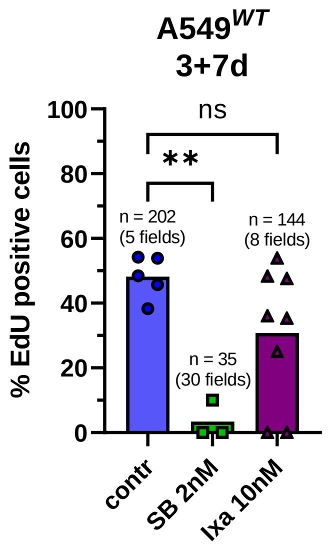


**Supplementary Figure S7.** Comparison of number of EdU-positive proliferating cells based on quantification of microscopy photos. A549$^{WT}$ treated with 2 nM SB743921 (SB) or 10nM Ixabepilone (Ixa) for 3 days with subsequent washout and 7 day incubation.

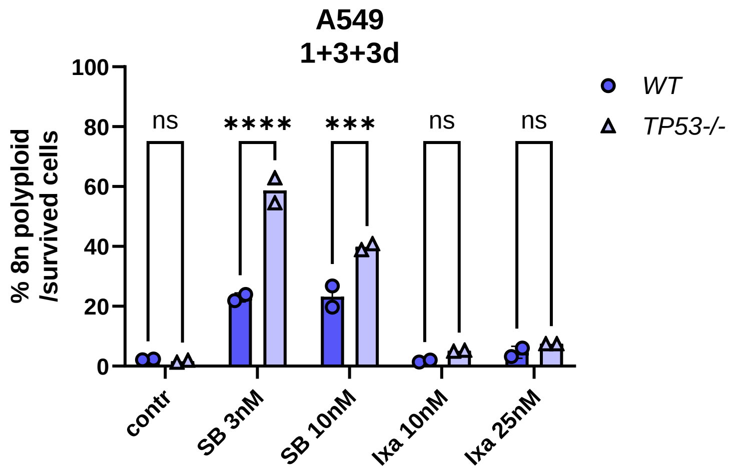


**Supplementary Figure 8.** Ratio of 8n polyploid cells to all survived cells. A549$^{WT}$ and A549$^{TP53-/-}$ cells treated with 3 and 10 nM SB743921 (SB) and 10nM and 25nM Ixabepilone (Ixa) for 1 day and subsequent 2 washouts every 3 days**.** Quantification based on flow cytometry data.

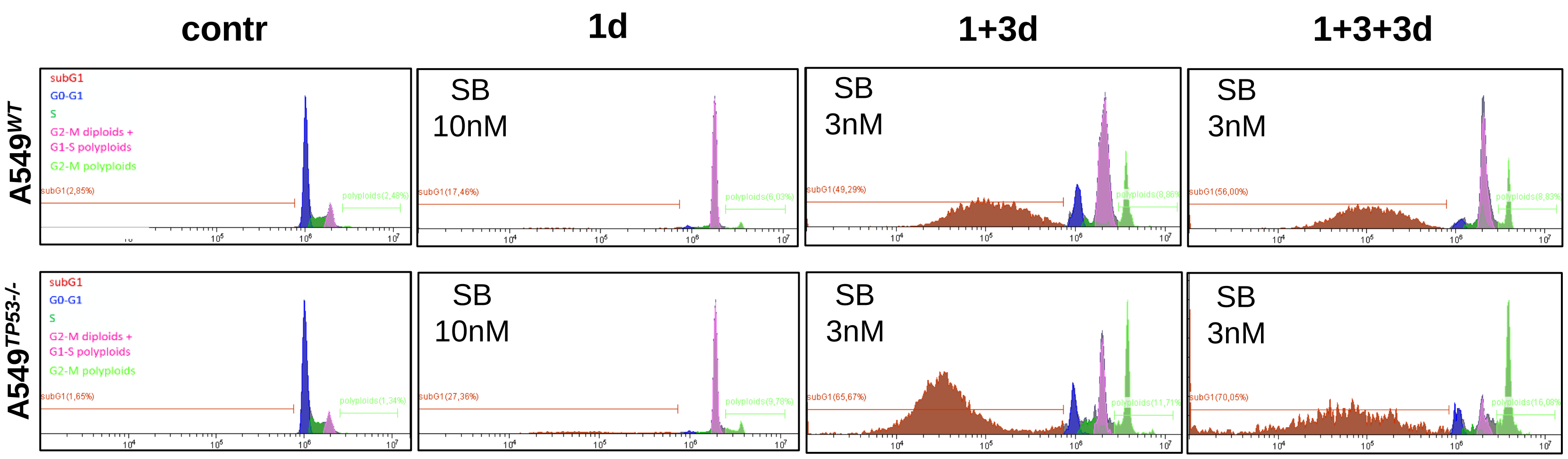


**Supplementary Figure S9.** Cell cycle distribution of A549$^{WT}$ (*upper panels*) and A549$^{TP53-/-}$ (*bottom panels*) treated with SB743921 (SB 10nM or 3nM) for 1 day and with or without subsequent washout(s). Flow cytometry data were obtained after incubation in PI-containing lysis buffer.

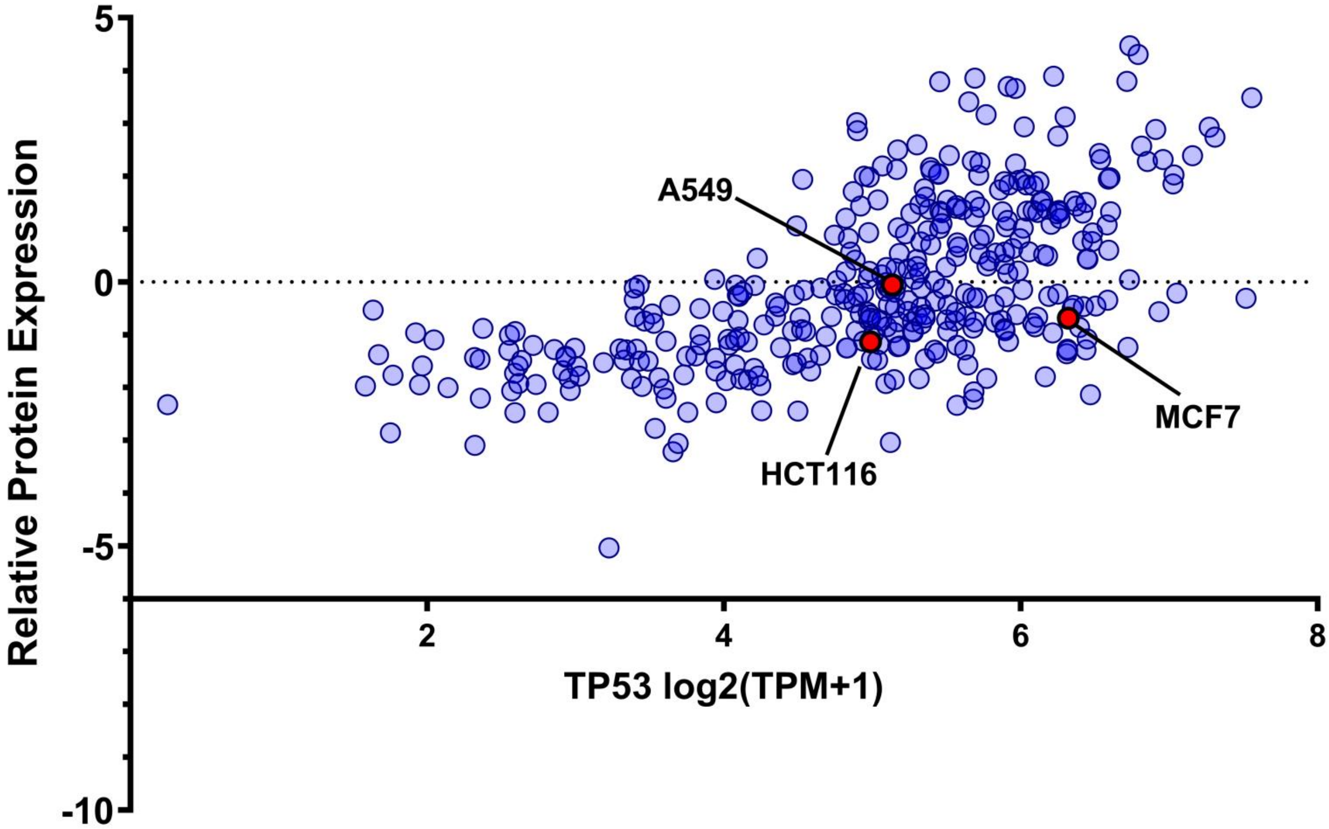


**Supplementary Figure S10.** Basic levels of *TP53* expression in A549, HCT116, MCF7 according to the DepMap database**.**
